# Higher Variability in Fungi Compared to Bacteria in the Foraging Honey Bee Gut

**DOI:** 10.1101/2020.10.20.348128

**Authors:** Leslie E. Decker, Priscilla A. San Juan, Magdalena L. Warren, Cory E. Duckworth, Cheng Gao, Tadashi Fukami

## Abstract

Microbial communities in the honey bee gut have emerged as a model system to understand the effects of host-associated microbes on animals and plants. The specific distribution patterns of bacterial associates among honey bee gut regions remains a key finding within the field. The mid- and hindgut of foraging bees house a deterministic set of core species that affect host health. In contrast, the crop, or honey stomach, contains a more diverse set of bacteria that is highly variable in composition among individual bees. Whether this contrast between the two gut regions also applies to fungi, another major group of gut-associated microbes, remains unclear despite their potential influence on host health. In honey bees caught foraging at four sites across the San Francisco Peninsula, we found that fungi were much less distinct in species composition between the crop and the mid- and hindgut than bacteria. Unlike bacteria, fungi were highly variable in composition throughout the gut, and much of this variation was attributable to bee collection site. These patterns suggest that the fungi may be passengers rather than functionally significant gut symbionts. However, many of the fungi we found in the bees have been recognized as plant pathogens. Assuming that some fungi remain viable after passage through the gut, the distribution patterns we report here point to the potential importance of honey bees as vectors of fungal pathogens and suggest a more prominent role of honey bees in plant pathogen transmission than generally thought.

**Importance (*Nontechnical explanation of why the work was undertaken*):** Along with bacteria, fungi make up a significant portion of animal- and plant-associated microbial communities. However, we have only begun to describe these fungi, much less examine their effects on most animals and plants. The honey bee, *Apis mellifera*, has emerged as a model system for studying host-associated microbes. Honey bees contain well-characterized bacteria specialized to inhabit different regions of the gut. Fungi also exist in the honey bee gut, but their composition and function remain largely undescribed. Here we show that, unlike bacteria, fungi vary substantially in species composition throughout the honey bee gut, contingent on where the bees are sampled. This observation suggests that fungi may be transient passengers and therefore unimportant as gut symbionts. However, our findings also indicate that honey bees could be major vectors of infectious plant diseases as many of the fungi we found in the honey bee gut are recognized as plant pathogens.

---

Gut microbes associated with the honey bee, *Apis mellifera*, have emerged as a powerful experimental system with which to uncover basic principles governing the assembly of host-associated microbial communities and their effects on host health (1). Gut microbes can influence host health by modifying carbohydrate metabolism, protein degradation, and resistance to parasites (2, 3). Because each region of the honey bee gut houses distinct microbial communities, different gut regions, including the crop, the midgut, and the hindgut, must be examined separately to understand the functional contribution and assembly rules of each community (4, 5).

For example, in the mid- and hindgut (hereafter intestine) of the honeybee, a deterministic set of functionally indispensable core microbes appear to exist across all healthy workers regardless of location (4, 6, 7). In contrast, the crop shows high heterogeneity in microbial species composition even among healthy workers, probably reflecting the spatial and temporal variation of ingested environmental microbes (8, 9) with potential consequences for host resistance to pathogens (10, 11). Critically, this research has focused almost exclusively on bacteria. It remains unknown whether the contrast between crop and intestinal communities applies only to bacteria or is also observed in other understudied groups of microbes, such as fungi, which may affect host health in ways that are currently underappreciated (12, 13).

In this study, we examined both bacteria and fungi in foraging workers to test two hypotheses: (1) fungal species composition is as distinct between the crop and the intestine as is bacterial species composition, and (2) fungi, like bacteria, show more variable species composition in the crop than in the intestine. To this end, we collected 101 *A. mellifera* foraging workers at four sites on the San Francisco Peninsula in California, USA (Fig.1d, Table S1). We dissected the entire gut, separating the crop from the intestine. We then extracted and sequenced the bacterial V4 region of the 16S ribosomal gene (505-806) and the fungal ITS1f-ITS2 region (14) (see Supplementary Information). Sequences were clustered into OTUs using VSEARCH (15), and taxonomy assigned for bacterial and fungal OTUs using QIIME (16) and UNITE (17).

**Figure 1.**
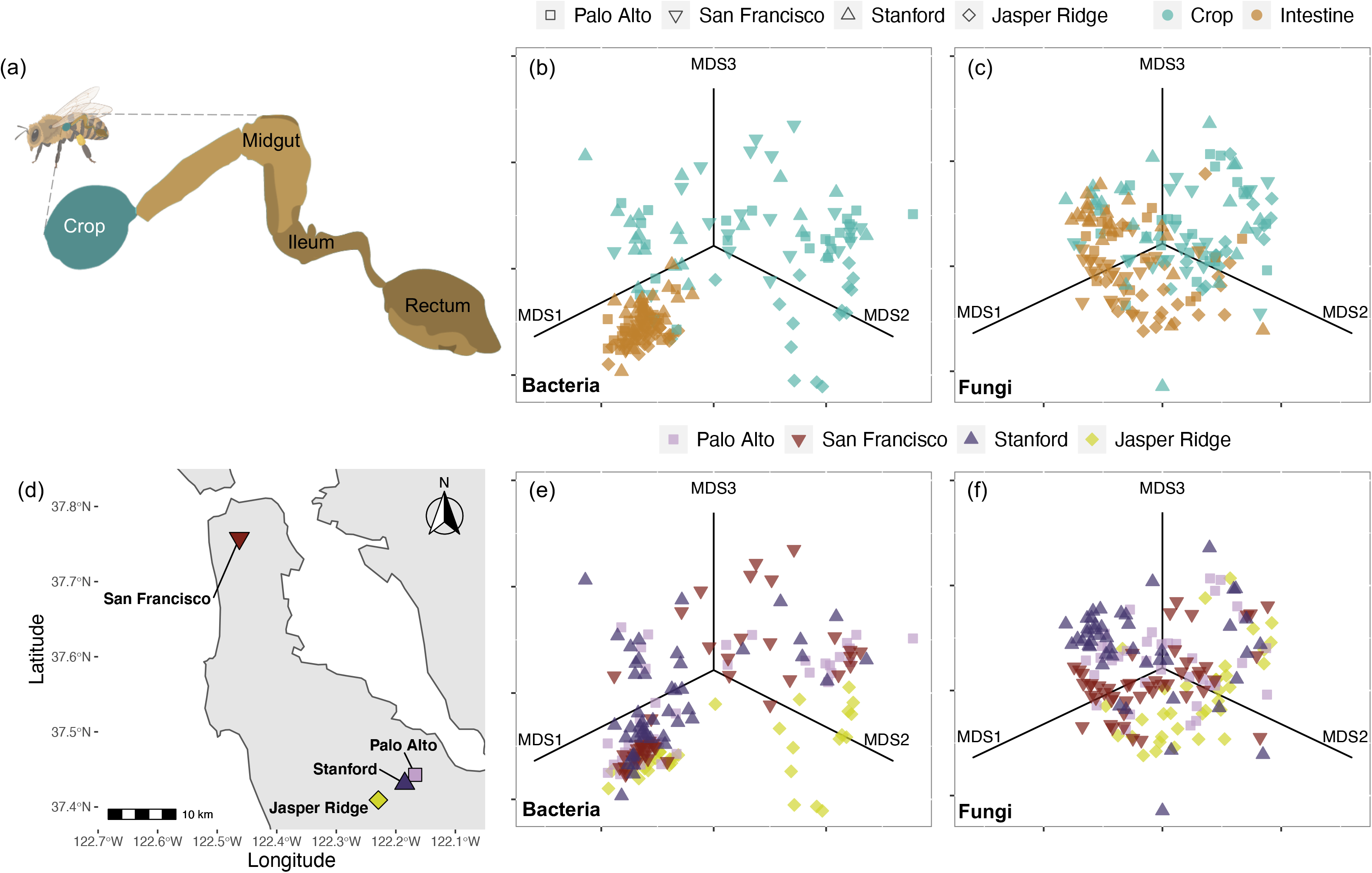
(a) Honeybee gut anatomy color-coded by sampling scheme dividing the gut into the crop (turquoise) and the remaining posterior intestine regions (gold). Non-metric multidimensional scaling (NMDS) based on Bray-Curtis dissimilarity matrices calculated from rarefied OTU tables show variation in (b) bacterial and (c) fungal community structure, where shape indicates collection site. Each dot indicates a bee individual. A map of the collection sites within the San Francisco Bay Area: Jasper Ridge Biological Preserve (N=25 bees), Stanford University (N=26 bees), Palo Alto (N=24 bees), and San Francisco (N=26 bees). (d). The same NMDS as in (b) and (c), but color-coded instead by collection site, illustrates weak retention of site effects in (e) bacteria and stronger site effects in (f) fungi.

As expected, bacterial community composition was most strongly predicted by gut region (PERMANOVA, gut region: R^2^=0.32, p=0.001, Fig.1b-c), and higher among-host variation was detected in the crop than in the intestine (Betadeviation(18): F_1,168_=14.5, p=0.0002). Although fungi also showed high variability in the crop (Fig. 1, Betadeviation: F_3, 161_=40.8, p < 0.0001, Fig.S1), fungi in the intestine were more diverse in species composition than bacteria (Shannon: F_4,330_=7.60, p=0.006, Fig.S2). Moreover, fungi retained more of the among-site differences from the crop to the intestine than did bacteria (Fig.1). Sample site was the strongest predictor of fungal species composition not just in the crop, but also in the intestine (PERMANOVA, site: R^2^=0.14, p=0.001, Fig.1e-f), with gut region explaining only a small proportion of fungal composition (PERMANOVA, gut region: site, R^2^=0.03, p=0.001).

To further quantify differences between bacteria and fungi, we applied the CLAM test, a multinomial species classification method (19), which differentiated OTUs into four categories: crop-associated, intestine-associated, generalist, or too rare to classify. We found that only 1% of bacterial OTUs were categorized as generalists, whereas 7% of fungal OTUs fell into this category (Fig.2a-d). Furthermore, only 2% of bacterial OTUs were classified as intestine-associated, whereas 11% of fungal OTUs were classified as intestine-associated.

**Figure 2.**
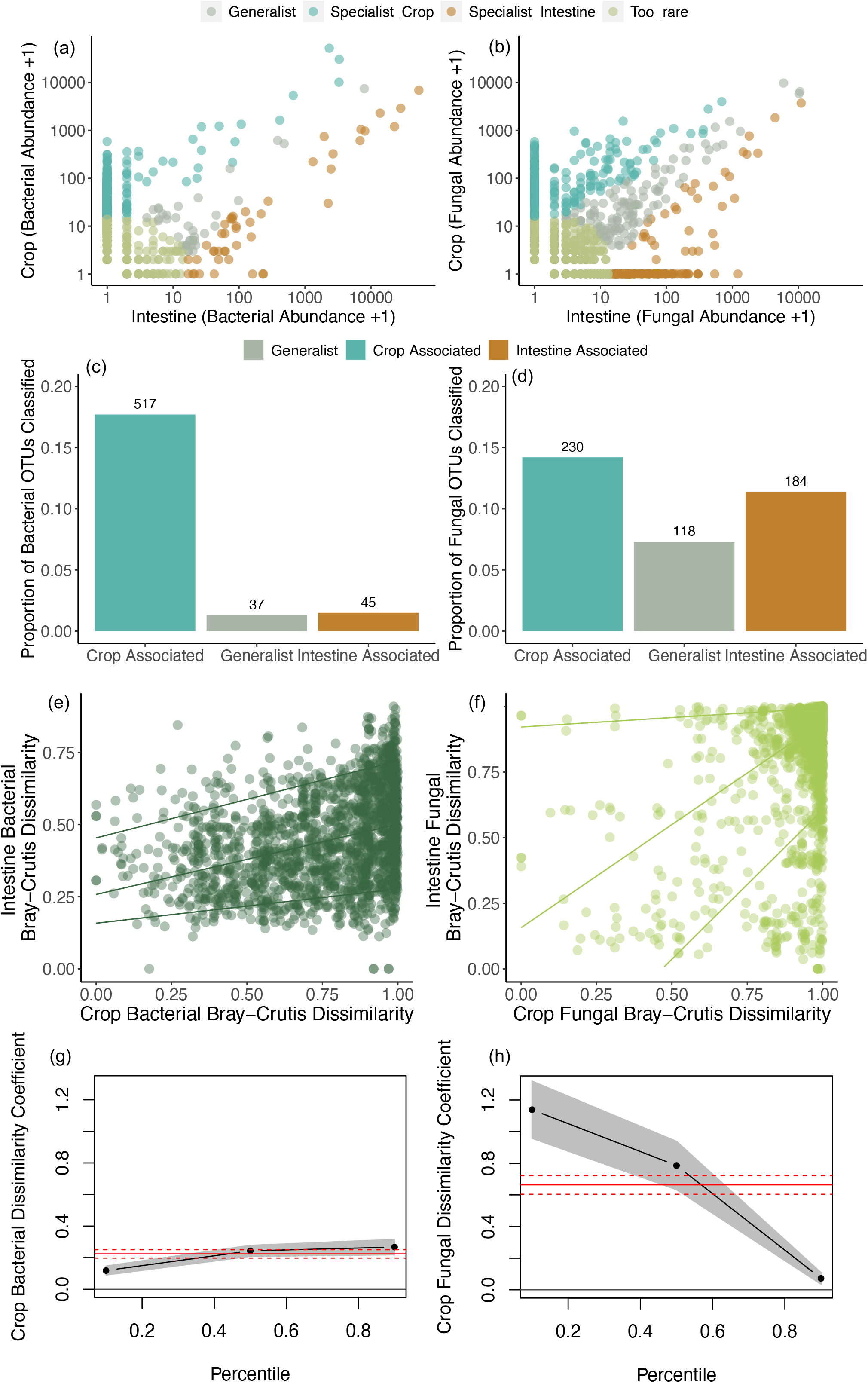
Fewer bacterial OTUs were categorized as intestine-associated specialists and generalists compared to fungi using a multinomial species classification method (clamtest) to sort OTUs into categories based on relative abundance in each domain: (a) bacteria and (b) fungi. Summarized classification results for (c) bacteria and (d) fungi. Relationships between (e) bacterial crop and intestine community structure (dark green), and (f) fungal crop and intestine community structure (light green). Lines are regressions against the 10^th^, 50^th^, and 90^th^ quantiles respectively from bottom to top. Changes in slope reflect the extent to which the composition of intestine communities depend on crop communities. Slope coefficients with 95% confidence intervals for each quantile are plotted for (g) bacteria and (h) fungi in relation to the ordinary least square regression coefficient plotted with 95% confidence intervals in red.

In addition, to determine how tightly crop composition was correlated with intestinal composition, we used Mantel tests to compare dissimilarity matrices between the crop and the intestine for bacteria and fungi separately. Crop composition was positively correlated with intestinal composition in both bacteria (Mantel r=0.34, p<0.0001, Fig.2e) and fungi (Mantel r=0.35, p=<0.0001, Fig.2f). The behavior of the fungi appeared nonnormal (Fig.2f), and we used quantile regression to determine how the composition relationships detected by the Mantel tests differed as dissimilarity values increased in magnitude from the mean. Bacteria in the intestine were correlated with those in the crop similarly across all three quantiles (10^th^ slope= 0.12, 50^th^ slope=0.24, 90^th^ slope=0.27, Fig.2g), whereas in fungi, the slope of the relationship depended on the quantile examined (10^th^ slope= 1.14, 50^th^ slope=0.79, 90^th^ slope=0.07, Fig.2h). Lastly, we calculated checkerboard scores describing species co-occurrence (20), which indicated significant segregation only in the crop in bacteria, but this pattern was not observed in fungi (Table S2).

Taken together, our results reject both of the hypotheses we set out to test, highlighting contrasting compositional patterns between bacteria and fungi in the honey bee gut. Specifically, we found that honey bees retained more of the across-site differences from the crop to the intestine in fungi than in bacteria. Furthermore, unlike the constrained set of bacterial species in the intestine (4), fungal species composition was highly variable not just in the crop, but also in the intestine.

The broad distribution of fungal taxa we found throughout the gut suggests that these microbes may simply be ingested from external sources and then excreted. Moreover, our Mantel test results indicate that various fungal taxa disappear in a seemingly stochastic fashion as they move from the crop to the intestine. These results contrast the deterministic filtering of bacteria from the crop to the intestine that has been documented previously (4, 6) and corroborated here. The deterministic filtering in bacteria is also indicated by the disappearance of a significant checkerboard pattern of segregation from the crop to the intestine in bacteria, a pattern that we observed only in bacteria and not in fungi (Table S2).

Many fungal OTUs that could be identified to the species level in our study were reported previously as plant pathogens rather than honey bee symbionts (Table S3). Assuming that some of these fungi remain viable as they pass through the gut (21), our study suggests a more prominent role of honey bees as vectors of plant fungal pathogens across landscapes than previously recognized. Transmission of phytopathogens accumulating on the surface of honey bees has already been implicated in the spread of bacterial and fungal pathogens (22, 23), but the extent to which fecal transmission of fungal pathogens contributes to plant epidemics remains unknown. Honey bee hives are often transported among multiple orchards and farms for pollination (24). In our study, the composition of gut fungal communities was specific to foraging sites. However, if honey bees act as vectors of plant-pathogenic fungi, those fungal pathogens that would otherwise be locally restricted might be transmitted more broadly when hives are transported.

In summary, we provide evidence that fungal species composition is not nearly as distinct between the crop and intestine as in bacteria, and that fungal species composition is highly variable across the entire gut, unlike bacteria. These findings suggest that most fungi found in the honeybee gut may be transient passengers rather than symbionts that affect the health of the host. However, these fungi may still be functionally important as bee-transported plant pathogens. Further studies focused on the relationship between bees and gut fungi may increase our understanding of plant disease transmission and pollination.

## Data availability

Upon acceptance, the 16s rRNA and ITS gene amplicon data sets generated by this study will be deposited and available in the Sequence Read Archives of the National Center for Biotechnology Information (NCBI).

## Supplemental Material

Supplemental material is available online only.

TEXT, DOCX file, 51 KB
FIG S1, PDF file, 146 KB
FIG S2, PDF file, 143 KB
FIG S3, PDF file, 139 KB
FIG S4, PDF file, 95 KB
TABLE S1, PDF file, 60 KB
TABLE S2, PDF file, 53 KB
TABLE S3, PDF file, 66 KB
TABLE S4, PDF file, 55 KB
TABLE S5, PDF file, 35 KB

## Acknowledgements

We thank Megan Morris for advice on DNA extraction, sequencing, and data analysis. We also thank Callie Chappell, Nick Hendershot, Jesse Miller, and Chih-Fu Yeh for comments. This work was funded by NSF (DEB 1737758). CED was supported by the Stanford Summer Research Program.

The authors declare no conflict of interest.

LED, PAS, CED, MLW, TF designed the study, and LED, PAS, CED, MLW collected samples. LED & CG analyzed data, and LED wrote the first draft of the manuscript. All authors contributed to editing the manuscript.

